# Evaluation of Plant- and Microbial-Derived Protein Hydrolysates as Substitutes for Fetal Bovine Serum in Cultivated Seafood Cell Culture Media

**DOI:** 10.1101/2024.03.27.587063

**Authors:** Arian Amirvaresi, Reza Ovissipour

## Abstract

This study seeks to explore alternatives by substituting or reducing the conventional 10% serum concentration in Zebrafish embryonic stem cell (ESC) growth media with protein hydrolysates sourced from peas, mushrooms, yeast, and algae. Notably, algae exhibited the highest protein content, optimal amino acid balance, and favorable functional properties. When applied at concentrations ranging from 1 to 10 mg/mL, all protein hydrolysates demonstrated pro-apoptotic effects and inhibited cell growth, particularly when used in conjunction with 10% serum. However, concentrations ranging from 0.001 to 0.1 mg/mL displayed anti-apoptotic properties and promoted cell proliferation. The study found that media containing 1% or 2.5% serum, along with 0.01 mg/mL of protein hydrolysates, supported cell growth effectively. Lactate Dehydrogenase (LDH) Activity served as an indicator of cell health and integrity under the specified conditions of protein hydrolysate supplementation. Cells cultured in serum-free media exhibited significantly decreased cell membrane integrity (*P < 0*.*05*) compared to those in regular media or media containing low serum (1% and 2.5%) along with low concentrations (0.01 mg/mL) of protein hydrolysates. Furthermore, analysis of Greenhouse Gas emissions (GHG) suggested that media formulations containing 1% serum combined with low concentrations of protein hydrolysates present a sustainable approach for cell-based seafood production.

## 1. Introduction

By 2050, the world population is projected to reach 10 billion, posing a significant challenge in meeting the global demand for food. According to the Food and Agriculture Organization (FAO), a 70% increase in meat production is required to address this growing need (FAO, 2020). Among the various animal-based meats, aquatic food products play a crucial role as an essential commodity, constituting approximately 20% of the animal protein consumed globally.

Cultivated meat involves the cultivation of cells or tissues in a carefully formulated medium that supports cell proliferation, metabolism, and differentiation. This innovative technique provides a promising solution to the challenges associated with traditional meat production. Cultivated meat reduces reliance on fisheries and aquaculture, mitigating environmental impacts, addressing animal welfare concerns, and reducing public health risks. It offers a sustainable and ethical solution to meeting global protein demand while overcoming challenges in traditional aquatic food production.

A culture medium plays a vital role in supporting the proliferation and growth of cells or tissues in the process of cultivated meat production (Yao and Asayama, 2017). The cost of cell culture media constitutes a significant portion of the total expenses involved in cultivated meat production, accounting for more than 99% of the total cost (Stout et al., 2022). One potential approach to reduce the overall cost of cultivated meat is to develop less expensive culture media. Therefore, finding alternative, cost-effective sources or formulations for these components could have a substantial impact on reducing the cost of cultivated meat production.

Fetal bovine serum (FBS) plays a crucial role in cell culture as it provides essential nutrients, growth factors, and hormones necessary for the proliferation and viability of cells. It has long been the gold standard in cell culture media due to its ability to support robust cell growth and differentiation (Stout et al., 2022). However, using FBS arouses several challenges due to its high cost presenting a significant barrier, impacting the scalability and commercial viability of cultivated meat production. Additionally, concerns about the ethical sourcing of FBS and the potential risk of disease transmission have prompted researchers to explore alternative, sustainable, and cost-effective options (Freshney, 2015). Developing suitable substitutes for FBS is essential to overcome these challenges and advance the field of cultivated meat towards a more scalable, affordable, and ethically sustainable future.

Protein hydrolysates, derived from protein cleavage, can be used as supplements in cell culture media to provide essential nutrients and growth factors for cells. Protein hydrolysates derived from various plant and animal sources have been utilized by researchers for cell growth and cultivation purposes (Batish et al., 2022). Peptides, derived from protein hydrolysates or synthesized chemically, play a role in cell signaling and can be incorporated into media to modulate cellular processes. The application of these peptides has gained significant attention due to their ability to provide essential nutrients and growth factors required for cell proliferation and maintenance. Researchers have explored the use of soy-derived peptides, among others, to support the growth of different cell types *in vitro* (Andreassen et al., 2022). These peptides offer a rich source of amino acids, which serve as building blocks for protein synthesis and cellular functions (Burteau et al., 2003; Spearman et al., 2014; Batish et al., 2022; Timoneda et al., 2024). Serum-free media can benefit from peptides in terms of cell growth, cell performance, differentiation and sustainability (Kim et al., 2011; Logarušić et al., 2021; Nikkhah et al., 2023; van der Valk et al., 2010).While, incorporating protein hydrolysates into the cell culture media, enhanced cell growth, proliferation and viability in many researches, the negative and pro-apoptosis properties of protein hydrolysates have been reported when applied at high concentrations between 1 to 10 mg/mL (Lee et al., 2008; Batish et al., 2022; Hsieh et al., 2022). In addition, sustainability and greenhouse gases (GHG) emissions have not been used to justify the application of protein hydrolysates in cell culture media. Thus, this research aims to evaluate the impact of different concentrations of serum, alone and in conjunction with different concentrations of various protein hydrolysates on fish cell line growth, morphology, viability, and performance. GHG emissions also is provided for different conditions.

## 2. Materials and methods

### 2.1. Materials and chemicals

The algae and pea protein were provided from Mountain Rose Herbs (Eugene, Oregon, USA) and NorCals organic (Crescent City, CA, USA), respectively. The white mushrooms (*Agaricus bisporus*) were obtained from the local market as fresh. Yeast hydrolysate, Bacto™ TC Yeastolate, serum, Lebovitz 15 media, HEPES and sodium bicarbonate were acquired from Thermo-Fisher scientific (Waltham, MA, USA). Alcalase^®^, a commercially available endoprotease enzyme (2.4 AU/g) derived from *Bacillus licheniformis* was obtained from Sigma-Aldrich Inc. (St. Louis, MO, USA). The Zebrafish Embryonic Stem Cells (ZESCs), ZEM2S CRL-2147^™^, were obtained from the American Type Culture Collection (ATCC). The antibiotics which were initially used in cell culture were obtained from Cytiva (Marlborough, MA, USA).

### 2.2. Protein hydrolysates production

The hydrolysates were prepared in at least six replicates, based on the optimum conditions (Table 1). The enzymatic hydrolysis of different substrates was conducted with minor adjustments tailored to each specific substrate, as described before (Batish et al., 2022, 2020). Briefly, the raw materials were homogeneously mixed with water at a predetermined ratio as specified in Table 1. Subsequently, the mixture was subjected to enzymatic hydrolysis at 60°C for 4 h. To stop the enzymatic reaction, the samples were subjected to 90°C for 10 min. The samples were centrifuged to collect the protein hydrolysates, which were then freeze-dried. The dry yield (%) was determined using equations 1. The lyophilized protein hydrolysates were stored at −20°C until further use.

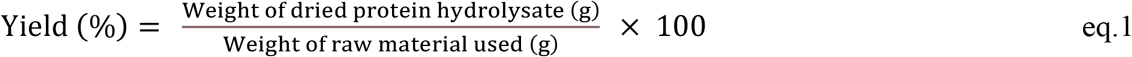

**Table 1.** Specific reaction conditions for enzymatic hydrolysis of substrates and the dry yield.

| Substrate | Water to substrate<br>ratio (v/v) | Enzyme<br>(%) | Hydrolysis<br>time (h) | Dry yield<br>(%) |
| --- | --- | --- | --- | --- |
| Algae | 10:1 | 2% | 4 | 23.8 ± 1.48 |
| Pea | 20:1 | 4% | 4 | 39.58 ± 1.74 |
| Mushroom | 10:1 | 2.5% | 4 | 5.78 ± 0.35 |

### 2.3. Amino acid analysis, protein content and degree of hydrolysis

The total amino acid analysis was performed following the guidelines specified in the Association of Official Agricultural Chemists (AOAC) 982.30 E (a,b,c) (AOAC, 2006). Upon complete sample digestion with 6N HCl, an ion exchange chromatography method with post-column ninhydrin derivatization and quantitation was employed. The determination of crude protein content was carried out using the AOAC standard method, Kjeldahl. To calculate the protein content, the crude nitrogen content was multiplied by the appropriate nitrogen to protein conversion factor (*Kp*) specific to each sample type. For algae, a *Kp* value of 6.35 (Safi et al., 2013), for pea, a *Kp* value of 5.36 (Mariotti et al., 2008), and for mushroom, a *Kp* value of 4.7 were applied (Mattila et al., 2002).

The protein quality assessment was conducted using Digestible Indispensable Amino Acid Score (DIAAS) as recommended by FAO (FAO, 2011) using equation 2:

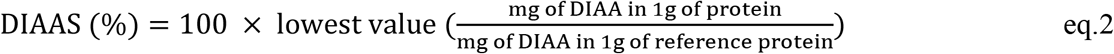

Degree of hydrolysis was measured using formal titration method (Taylor, 1957).

### 2.4. Oil holding capacity (OHC)

The evaluation of the oil holding capacity (OHC) was carried out for each protein hydrolysate (Shahidi et al., 1995). Briefly, 500 mg of protein hydrolysate was combined with 10 mL of pure canola oil. The mixture was then left at room temperature for 30 min, gentle mixing every 10 min. Subsequently, centrifugation was performed at 2500*g* for 10 min at room temperature. The OHC was determined by utilizing equation 3.

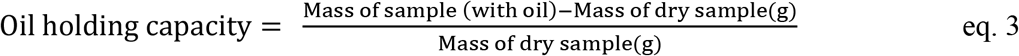

### 2.5. Emulsifying capacity (EC)

The evaluation of the EC involved the mixing of 500 mg of protein hydrolysates with 50 mL of a 0.1 M sodium chloride solution in a 250 mL conical flask, conducted at ambient temperature (Yasumatsu et al., 1972). The EC was determined by employing equation 4.

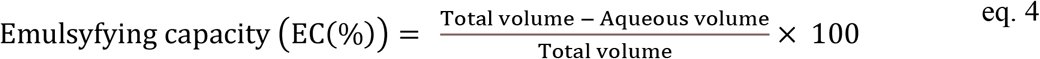

### 2.6. Foaming capacity (FC)

The foaming capacity (FC) was determined based on the methodology described by Pacheco-Aguilar et al. (2008). In this procedure, 750 mg of protein hydrolysates were mixed with 25 mL of distilled water for a duration of 10 minutes. Subsequently, the mixture was subjected to homogenization for two minutes. The FC was then calculated utilizing equation 5.

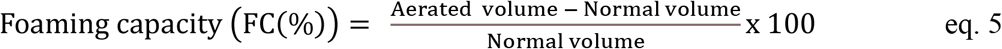

### 2.7. Cell culture and maintenance

For the initial cell culture, a combination of Lebovitz-L-15 media (L-15), Dulbecco’s Modified Eagle Media (DMEM), and Ham’s F12 Media (F-12 media) was utilized in a ratio of 15:50:35, respectively. The culture media were supplemented with buffering agents, including 20 mM 4-(2-hydroxyethyl)-1-piperazineethanesulfonic acid (HEPES) and 0.18 g/L sodium bicarbonate, and supplemented with 10% FBS. The cells in ampoules were thawed at a temperature of 28°C and subsequently resuspended in 9 mL of media containing 5% fetal FBS within a T-25 cm^2^ flask. The cells were allowed to settle for 30 min, following which additional FBS was added to achieve a final concentration of 10% FBS. The subculturing procedure was conducted when the cells reached 80-85% confluence. This was accomplished by rinsing the cells with phosphate-buffered saline (PBS) and subsequently treating them with Tryple Express, a cell detachment reagent, to detach the cells from the surface of the flask. The flask containing the detached cells was maintained at a temperature of 28°C for a duration of 5 minutes to ensure complete detachment. Subsequently, the cells were transferred to a 15 mL Falcon tube. To neutralize the Tryple reagent, serum-free media (L-15 media) was used, and the cell suspension was then subjected to centrifugation at 130*g* for 8.5 minutes. The supernatant was carefully discarded, and the resulting cell pellet was resuspended in media containing 10% serum. The cell number and viability were assessed using an automatic cell counter, and cell splitting was performed based on the obtained results. The cells reached confluency within one week, with a media change performed once on the third day.

### 2.8. Cell performance in reduced or serum free media

To evaluate the potential capability of protein hydrolysates in reducing serum usage, various protein hydrolysates were applied at different concentrations in two different experiments. First in Experiment I, the impact of different concentrations of serum (0, 1, 2.5, 5, and 10%) was evaluated on the cell performance. In Experiment II, various concentrations of protein hydrolysates (0.001, 0.01, 0.1, 1, and 10 mg/mL) were studied in combination with media containing 0, 5, and 10% serum. In experiment III, lower concentrations of protein hydrolysates (0.001, 0.01, and 0.1 mg/mL) in combination with media containing lower concentrations of serum (0, 1, 2.5, and 10%) were studied. The protein hydrolysate samples were prepared by combining them with sterile water that had been filtered through a 0.22 µm filter. The cells were initially seeded at a density of 50,000 cells/mL and incubated for 24 hours at 28°C in a medium containing 10% serum to facilitate cell attachment to the culture plates.

Cell numbers and morphology were documented at 24-hour intervals for a total of three days using the CKX-CCSW Confluence Checker, which captured three images per well. Each hydrolysate condition was tested in four biological replicates, with each replicate having three technical replicates. The growth rate was determined using equation 6, while the doubling time was calculated using equation 7.

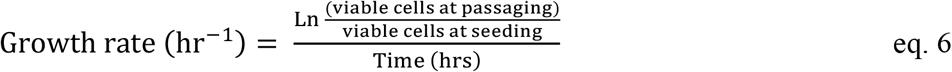

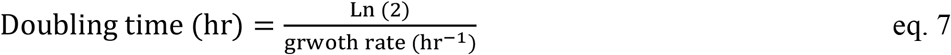

Cell viability was assessed utilizing PrestoBlue analysis. After 72 hours of cell seeding, 100 µL of PrestoBlue reagent was added to the wells, and the plate was incubated at 28°C for 2 hours. Subsequently, the absorbances at 570 nm and 600 nm were measured using a microplate reader. The reduction of the dye was determined by applying equation 8 for calculation.

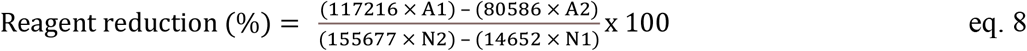

In the equation, A1 represents the absorbance of the sample wells at 570 nm, while A2 corresponds to the absorbance of the sample wells at 600 nm. Similarly, N1 denotes the absorbance of the media-only wells at 570 nm, and N2 represents the absorbance of the media-only wells at 600 nm.

### 2.9. Fluorescent imaging

Based on the results obtained from cell growth assays, optimal concentrations of protein hydrolysates were selected for further image analysis. To prepare the cells for imaging, they were first fixed with 4% paraformaldehyde for 10 min, followed by two rinses with PBS to remove the paraformaldehyde. To facilitate permeabilization, the cells were then incubated in 0.1% Tween solution for 10 min, followed by another round of rinsing with PBS. For visualizing the cell nuclei, Hoechst dye was diluted in PBS at a ratio of 1:2000, added to the cells, and allowed to incubate in the dark for 10 minutes. After incubation, the cells were washed twice with PBS, and 1 mL of PBS was added to each well. Subsequently, a second dye for visualizing the cytoskeleton, actin green dye, was added to each well, and the cells were incubated for 30 minutes before being washed twice with PBS. To optimize the image quality, the cells were then immersed in a live cell imaging solution. Finally, the cells were observed using a fluorescent microscope, with UV excitation/emission wavelengths of 361/486 nm for Hoechst dye and blue-cyan light with excitation/emission wavelengths of 495/518 nm for actin green dye.

### 2.10. Lactate dehydrogenase activity

The activity of lactate dehydrogenase (LDH) was assessed in different concentrations of selected protein hydrolysates supplemented with either 1% or 2.5% serum, which were found to be optimal for supporting cell growth in experiment III. The cells were cultured under these optimal conditions for a duration of three days. Following the three-day cultivation period, the supernatant was collected and transferred to 96-well plates for the LDH activity assay. The assay was performed according to the manufacturer’s instructions. To ensure statistical reproducibility, four biological replicates were included, with each biological replicate having three technical replicates.

### 2.11. Greenhouse gas emissions (GHG)

Greenhouse gas emissions associated with serum, intact proteins, and enzymatically hydrolyzed proteins were sourced from various inventories. The functional unit (FU) for this study is based on 1 kg of the products. Algae, mushroom, and pea were acquired as dried materials and were subjected to enzymatic hydrolysis in our laboratory. However, for yeast, protein hydrolysates were procured directly from the vendor. Thus, GHG emission data for yeast peptone (protein hydrolysates) was obtained from inventory. For algae, GHG emissions ranged from 1.5 to 150 kg CO_2_ eq./kg, contingent upon the production method and carbon sequestration data. Production method were obtained from the vendor website, and GHG emissions were selected based on a review of multiple articles (Clarens et al., 2010; Handler et al., 2012; Nette et al., 2016; “The Climate Intelligence Platform - CarbonCloud).

### 2.12. Statistical analysis

The statistical analysis was carried out using JMP 16 software. To ensure the validity of the analysis, the data was assessed for normality and homoscedasticity, confirming a normal distribution. Normality was tested using the Shapiro-Wilk test and Normal Quantile Plots (NQP), while the homogeneity of variability was assessed using Levene’s test. If the data exhibited a normal distribution, One-Way ANOVA and Tukey’s HSD (Honestly Significant Difference) tests were performed. However, if the data did not meet the criteria for normality, the Wilcoxon/Kruskal-Wallis test and Steel-Dwass/Wilcoxon pair comparisons were employed, along with their respective control conditions. Statistical significance was determined by considering differences with *P < 0.05*.

## 3. Results and discussion

### 3.1. Protein hydrolysates properties

For producing hydrolysates that properly support animal cell culture growth, the whole protein-rich substrate was used instead of protein isolate or concentrate. The yield of protein hydrolysates were examined to determine their potential for large-scale manufacturing (Table 1). As shown later in this investigation, the concentration of protein hydrolysate required for animal cellular culture applications is between 0.001 mg/mL to 0.1 mg/mL, and the yield of hydrolysates reported in this research appear to be suitable for cultivated meat scaling up. Producing cell-based meat requires the cultivation of billions of cells using limited space, time, and resources. One 2 L batch of any protein hydrolysate would be sufficient to supply the maximum capacity of a stirred bioreactor used for animal cell culture which has a capacity of 2000 L to produce 10 to 100 kg cell-based meat (Bellani et al., 2020).

### 3.2. Amino acid analysis, protein content and protein quality

The amino acid composition, protein content, DIAAS score and protein quality are presented in Table 2. All protein hydrolysates contained a high concentration of amino acids with cell-growth promoting properties, such as alanine, glycine, proline, and aspartic acid (Lu and Zhao, 2018). All protein hydrolysates examined in this study demonstrated a notable abundance of glutamic acid, a crucial component in animal cell culture. Glutamic acid serves as a significant nitrogen source for in vivo processes and contributes to the synthesis of amino acids, proteins, and nucleic acids. Furthermore, glutamic acid acts as a substantial supplier of metabolites in the tricarboxylic acid (TCA) cycle, supplying carbon and nitrogen for biosynthesis (Hosios et al., 2016). The hydrolysates were also found containing significant amounts of hydrophobic amino acids, including glycine, alanine, valine, and leucine. Furthermore, both yeast and algae protein hydrolysates exhibited a high protein quality, as evidenced by their DIAAS content of essential amino acids.

**Table 2.**
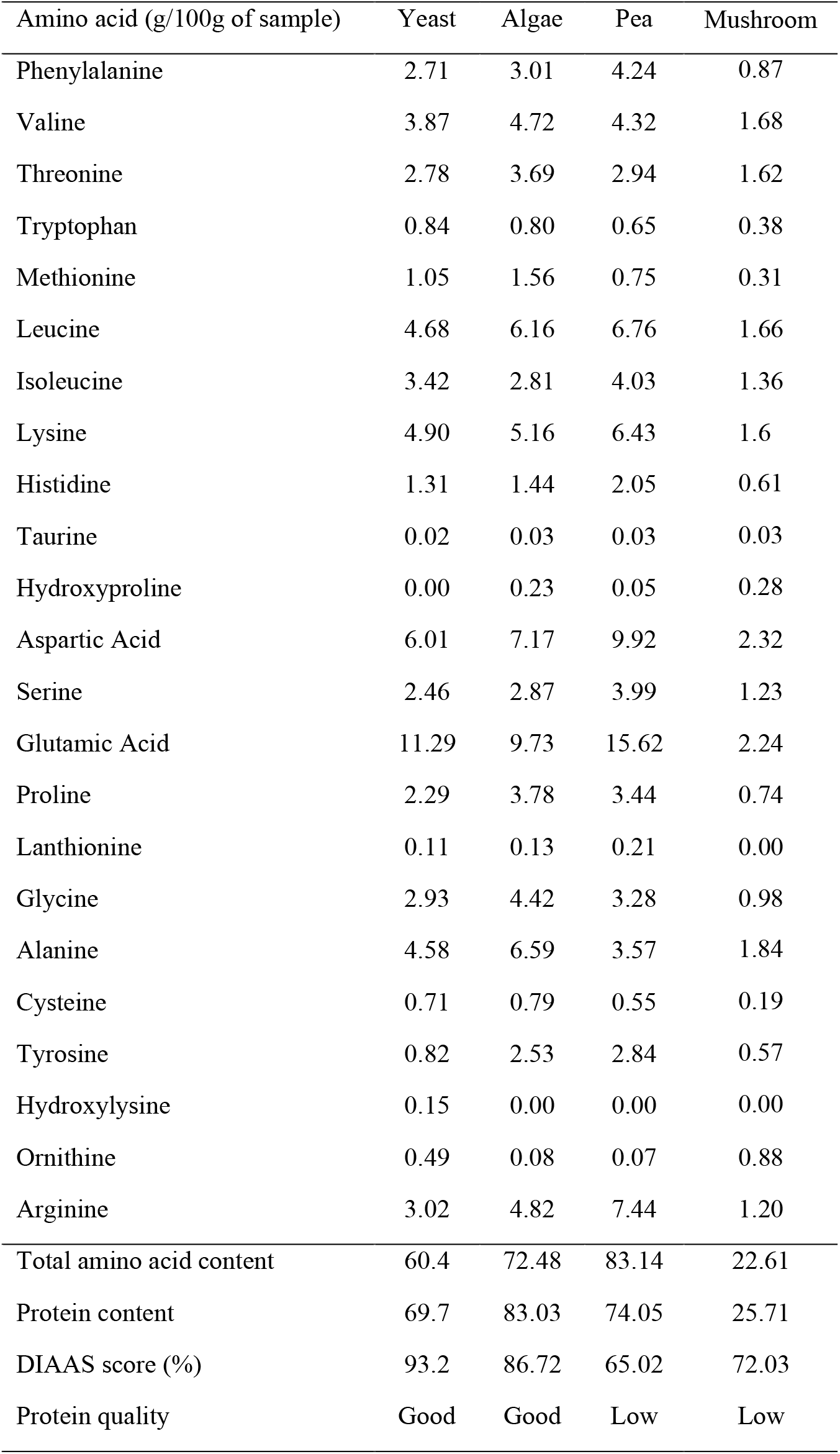
Amino acid and protein contents, DIAAS score, and quality of all hydrolysates.

### 3.3. Degree of hydrolysis and techno-functional properties

The results of the degree of hydrolysis and functional properties are presented in Fig. 1A-D. Degree of hydrolysis of were 15, 20, 40, and 46%, for pea, algae, yeast and mushroom, respectively. Yeast and mushroom hydrolysate showed the highest degree of hydrolysis, 40% and 46.1%, respectively. Degree of hydrolysis can influence the level of peptides releasing, the size, conformation, and amino acid sequence impacts various functional and bioactive properties to the hydrolysate product (Hall et al., 2017; Monaya et al., 2022).

**Fig. 1.**
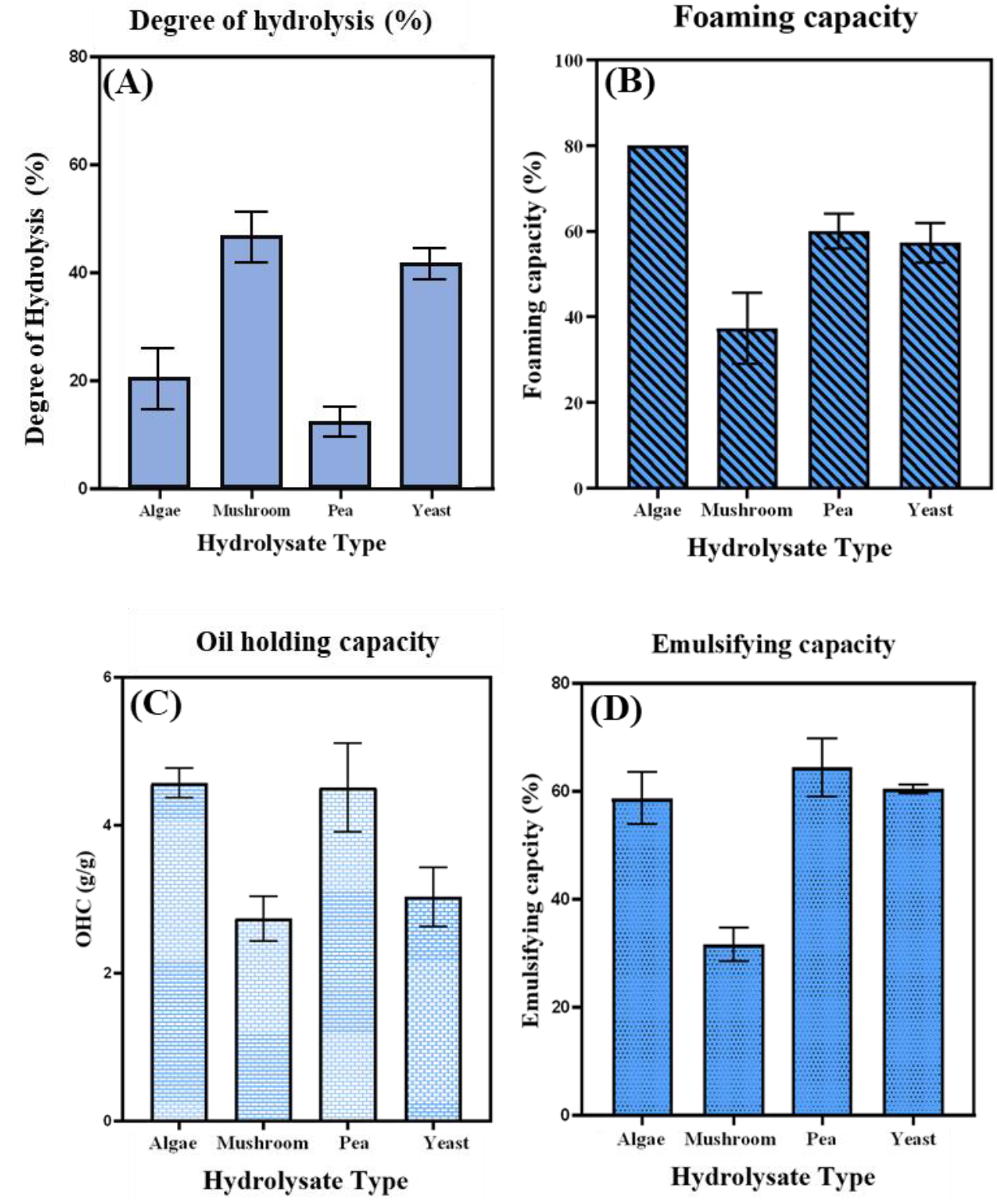
Techno-functional properties of protein hydrolysates. (A) Degree of hydrolysis; (B) foaming capacity; (C) Oil holding capacity; (D) emulsifying capacity.

Algae protein hydrolysates showed the highest foam capacity, followed by pea and yeast protein hydrolysates, while, mushroom protein hydrolysates showed the lowest foam capacity. Foaming properties are related to partially denaturing a protein and exposing as its hydrophobic regions. These regions can efficiently absorb air-water interfaces and form a lower interfacial tension, thus enhancing foaming capability. However, extensive denaturation will reduce foaming capability of the proteins (Mauer, 2003). Foaming in cell culture in bioreactor environment, may cause issues, such as inhibiting cell development by limiting the surface area contact between the growth media and the bioreactor headspace and decreasing oxygen transfer rates. Therefore, ideal hydrolysates are expected not to increase the foaming that occurs spontaneously in a bioreactor when added to media. The highest oil holding and emulsifying capacities were related to algae and pea protein hydrolysates, followed by yeast and mushroom hydrolysates.

### 3.4. Cell morphology, growth, and viability

In this study, protein hydrolysates from different sources were applied to reduce or fully replace serum. The following results demonstrate the changes in cellular morphology, growth, and viability of zebrafish embryonic stem cells growing under different media conditions with or with serum and at various concentrations of protein hydrolysates.

### 3.5. Impact of serum concentration

For culturing zebrafish ESC, 10% serum is the gold standard culture condition for proper growth of cells. In Experiment I, different concentrations of serum, 0, 1, 2.5, 5 and 10%, were evaluated for cell growth and doubling time. The impact of serum at various concentrations on the parameters of cell growth and viability is demonstrated in Fig. 2.

**Fig. 2.**
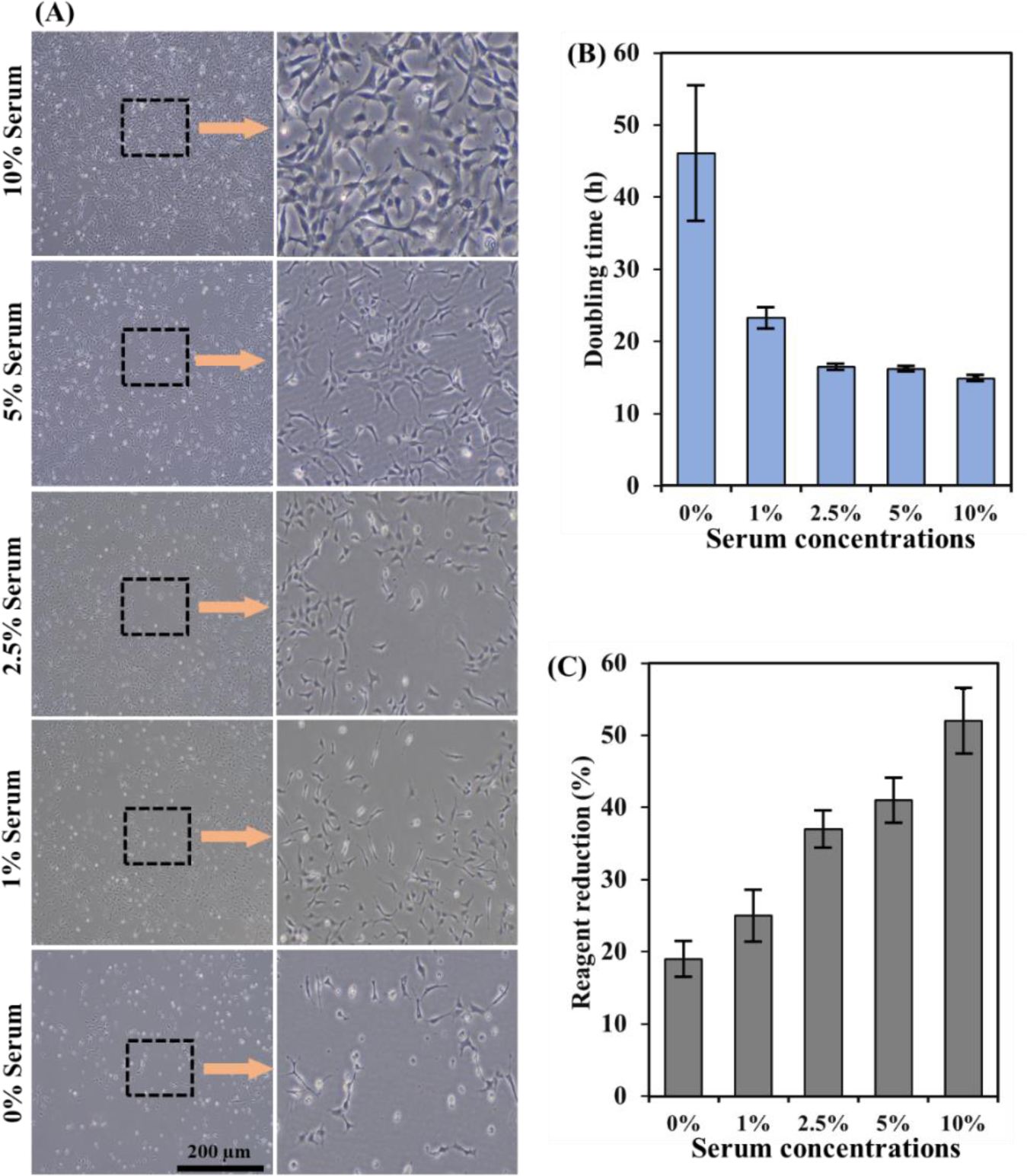
Cell growth parameters of ZEM2S cells at various serum concentrations. (A) micrographs of ZEM2S cells obtained on day 3; pictures in **the** right column are 20X enlarged from the inserted boxes, indicating the cell density and morphology, which significantly changes after removing serum; (B) doubling time indicating that completely removing serum will significantly increase the doubling time; (C) Cell viability results from PrestoBlue cell viability tests, indicating significantly higher cell activity in cells which were cultured in 10% serum.

After 24 hours of incubation in media containing 10 % serum, the test media were changed into media containing 0, 1, 2.5, 5, and 10% serum, then the growth of the cells was studied on a daily basis for three days using a phase-contrast microscope equipped with image analysis software. Doubling time was increased from 14.85 h in 10% serum media to 46 h after completely removing serum. As the serum concentration decreased, the cell density decreased significantly (*P < 0.05*). By decreasing the serum from 10% to 5%, while cell numbers reduced from 1.4 × 10^6^ to 1 × 10^6^ cell/mL, the cell morphology has not changed (Fig. 2A). At a serum-free media condition, there was a dramatic decrease in cell number from 1.4 × 10^6^ to 1.8 × 10^5^ cell/mL, and morphological changes with an increase in rounded cells indicating the cell death (Fig. 2). Complete serum removal causes cell starvation, apoptosis, and cell morphology changes which is in agreement with other researchers findings (Pirkmajer and Chibalin, 2011; Terra et al., 2011).

In Experiment II, we evaluated the impact of different protein hydrolysates concentrations from 0.001 to 10 mg/mL in combination with different concentrations of serum (0, 5, and 10%) on cells performance including cell growth (Fig. 3A, C, E, G, I) and cell viability (Fig. 3B, D, F, H, J). Overall, the cell density and cell viability results indicated that 1 and 10 mg/mL protein hydrolysates significantly reduced the cell growth and cell viability in media containing high concentrations of protein hydrolysates (*P < 0.05*). Serum-free media containing 10 mg/mL of mushroom and yeast hydrolysates, indicated higher cell growth compared to serum containing media supplemented with 10 mg/mL of protein hydrolysates. Providing protein hydrolysates at high concentrations for serum containing media (5 and 10%), also reduced cell viability and cell growth.

**Fig. 3.**
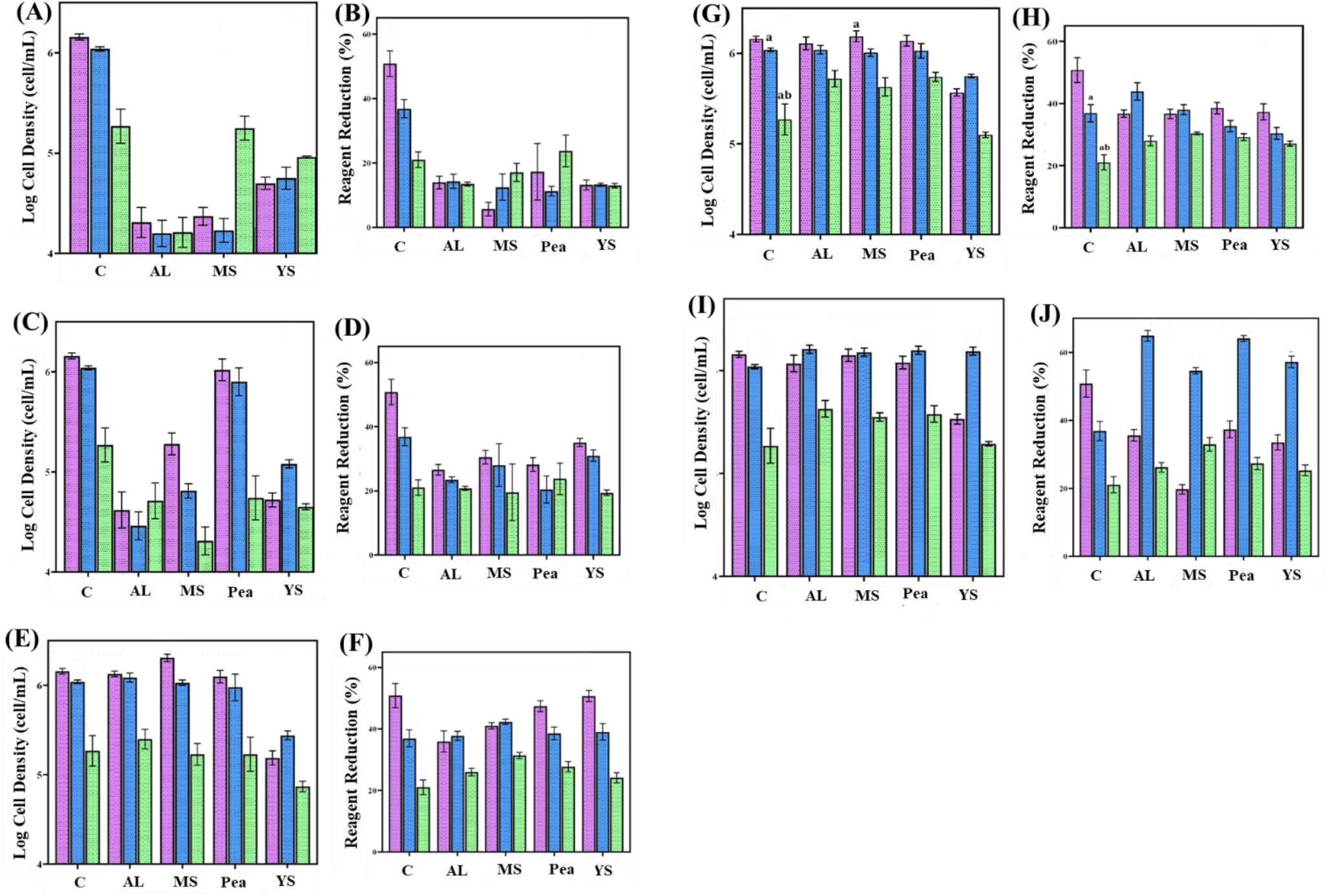
ZEM2S cells growth in media containing different serum concentrations (0, 5 and 10%) supplementing with various protein hydrolysates including C: Controls (no protein hydrolysates; AL: Algae; MS: Mushroom; Pea; YS: Yeast, at different concentrations including: (A, B) 10 mg/mL; (C, D) 1 mg/mL; (E, F) 0.1 mg/mL; (G, H) 0.01 mg/mL; (I, J) 0.001 mg/mL. Pink: 10% Serum; Blue: 5% serum; Green: 0% serum. Note: In experiments with 10 mg/mL protein, pea protein was not dissolved in media. Thus, the data was not collected for high concentrations of pea. Cell growth (A, C, E, G, I); Cell viability (B, D, F, H, J).

Within 24 hours of adding a medium containing high concentrations of protein hydrolysate, the cells lost their fibroblast-like morphology, and cell death was observed. In contrast, the effects of low concentrations of protein hydrolysates (0.001, 0,001, and 0.1 mg/mL) on cells growth, and viability illustrated that the lower concentrations of protein hydrolysates supported cell growth and viability (Fig. 3). At lower concentrations (0.001, 0.01 and 0.1 mg/mL) of protein hydrolysates, serum-free media containing protein hydrolysates improved cell growth and viability compared to the serum-free media control. The results suggested that algae, mushroom, and pea protein hydrolysates could potentially support cell growth in a serum-free environment. When serum was reduced to 5%, these protein hydrolysates successfully increased cell number more than the media containing 10% serum. Protein hydrolysates have been demonstrated to have more than a nutritional effect on cells, including protective, antiapoptotic, and growth-promoting properties. In Experiment III, the impacts of lower concentrations of protein hydrolysates (0.001, 0.01, and 0.1 mg/mL) in combination with low concentrations of serum (1 and 2.5%) on cell growth, viability (Fig. 4), and LDH activity (Fig. 5B) were evaluated.

**Fig. 4.**
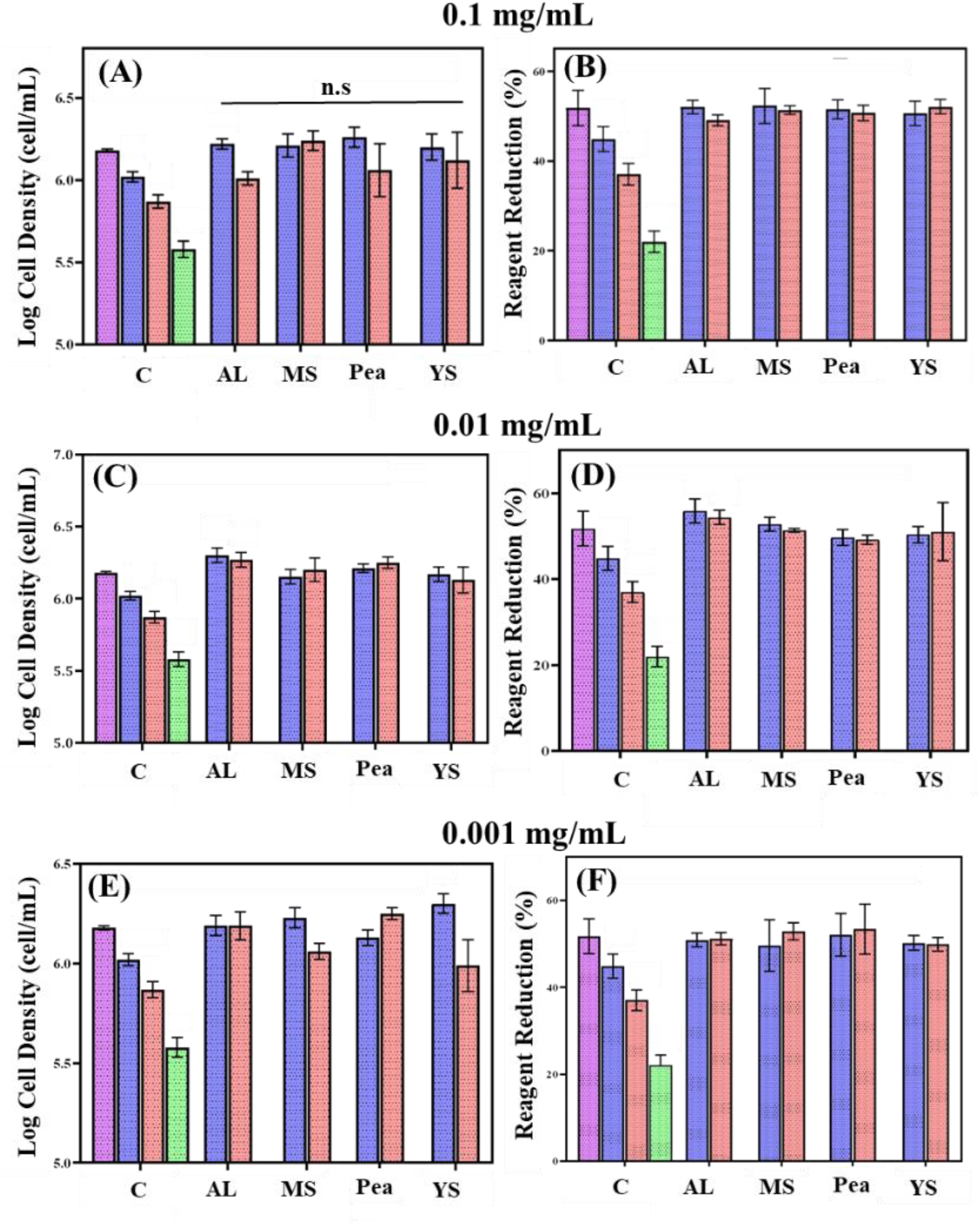
ZEM2S cells growth (A, C, E) and cell viability (B, D, F) in media containing different serum concentrations (0, 1, 2.5 and 10%) supplementing with various protein hydrolysates including C: Controls (no protein hydrolysates; AL: Algae; MS: Mushroom; Pea; YS: Yeast, at different concentrations including: 0.1 mg/mL; 0.01 mg/mL; 0.001 mg/mL. Pink: 10% Serum; Blue: 2.5% Serum; Orang: 1% Serum; Green: 0% serum.

**Fig. 5.**
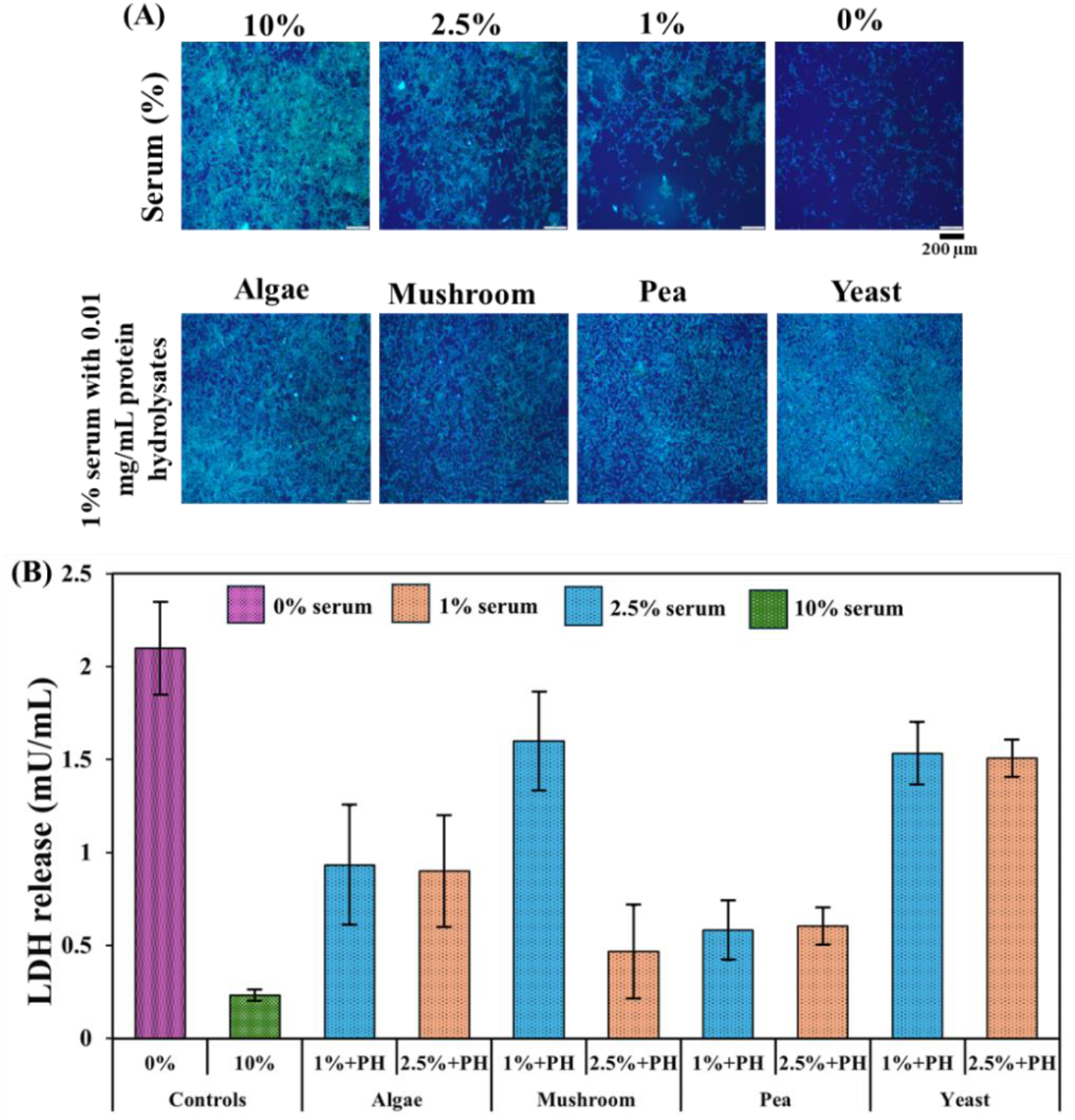
(A) Staining images from cells grown at different concentrations (0, 1, 2.5, and 10%) of serum, and cells grown only in 1% serum containing 0.01 mg/mL of different protein hydrolysates; (B) LDH activity for cells in serum-free media (0%), 10% serum, 1 and 2.5% serum supplemented with 0.01 mg/mL protein hydrolysates.

The results indicated that providing protein hydrolysates at 0.001, 0.01 and 0.01 mg/mL in combination with 1 and 2.5% serum, will increase cell growth and cell viability more than control groups as serum-free, and media containing 1, 2.5, and 10% serum. These results indicate that for fish cell lines such as zebrafish fibroblast cells, serum in media could be reduced up to 90% without compromising cell growth, and cell viability, when protein hydrolysates are provided at relatively low concentrations. No significant differences were observed among the protein hydrolysates in terms of the cell growth and cell viability.

A study using several plant peptones (protein hydrolysates) in CHO cell lines revealed that adding these hydrolysates did not increase the nutritive value of proteins, but dramatically boosted cell growth (Burteau et al., 2003). Similar effects were reported when CHO cells were grown with 10% serum and algal extracts (Ng et al., 2020). On CHO-K1 cell lines with cottonseed hydrolysate, this synergistic impact of protein hydrolysates with other medium components was demonstrated by increased cell biomass (Babcock and Antosh, 2012). In addition, providing rapeseed protein hydrolysates reduced cell death rate and increase culture time up to 160 h instead of 80 h in CHO cells (Farges-Haddani et al., 2006). Soy hydrolysates promoted CHO cells growth without increasing protein productivity, while, yeast hydrolysate reduced cell growth, but increased the level of protein productivity glycosylation profile (Spearman et al., 2014).

While, lower concentrations of protein hydrolysates support cell growth, higher concentrations, (4 to 6 mg/mL) of protein hydrolysates, including soy (Lee et al., 2008) and fish gelatin (Hsieh et al., 2022), significantly decreased cell proliferation. Our previous study results also indicated that oyster, mussel, lugworm, cricket, and black soldier fly larvae protein hydrolysates reduce cell proliferation at high (1 and 10 mg/mL) concentrations when combined with higher concentrations of serum (10%) (Batish et al., 2022; Timoneda et al., 2024). In addition, wheat gluten hydrolysate between 6 and 12 mg/mL reduced cell proliferation (Radošević et al., 2016). Protein hydrolysates derived from various wastes, such as eggshells and carcasses, at 10 mg/mL had a similar dose-dependent cytotoxic effect on bovine stem cells (Andreassen et al., 2020). This might be related to the nutrient balance due to high amino acid or oligopeptide concentrations (Chun et al., 2007; Hsieh et al., 2022; Zhang et al., 1994). On the other hand, low concentrations of whey protein hydrolysate, *i.e*. 0.01-0.5 mg/mL, boosted osteoblastic cell proliferation, viability, and alkaline phosphatase activity (Jo et al., 2020). Similar results were observed in bovine cell culture with algal extract and various other industrial byproduct hydrolysates; cell growth increased compared to serum-free conditions, however less than in serum-rich conditions (Andreassen et al., 2020). It has been shown that the cell growth in the presences of protein hydrolysates at different concentrations of serum depends on the degree of hydrolysis (Girón-Calle et al., 2010). However, in this study, no correlation was observed between degree of hydrolysis (15-46%, Fig. 1A) and cell growth. This could be explained by the fact that different sources of biomass were applied in this study, while, other researchers only used one specific biomass with different degree of hydrolysis (Girón-Calle et al., 2010).

Overall, protein hydrolysates from different sources could potentially enhance cell growth in low-serum or serum-free media. Protein hydrolysates provide several benefits for cells, due to their free amino acids, chemical compositions, peptides structures, bioactive properties such as anti-apoptotic effects (Spearman et al., 2014), anti-oxidation properties (Ovissipour et al., 2013), immunomodulatory effects, and antibacterial properties (Spearman et al., 2014). However, using protein hydrolysates may have several drawbacks including pro-apoptotic properties at higher concentrations, being undefined, leading to batch-to-batch quality challenges, being contaminated with other by-products from enzymatic hydrolysis process (Spearman et al., 2014). For example, microbial protein hydrolysates from yeast may contain specific peptides from the cell wall, or bacteria hydrolysates may have high level of lipopolysaccharide (LPS), an endotoxin which is difficult to completely remove from the final product (Lobo-Alfonso et al., 2010). In addition, accumulation of certain amino acids such as tryptophan in the culture, may inhibit cell growth (Alden et al., 2020). The rapidly growing cultivated meat industry requires innovative, sustainable, and cost-efficient media components. Protein hydrolysates sourced from affordable and eco-friendly biomass materials hold promise in bolstering this sector. Nonetheless, further investigation into emerging biomass conversion methods, the influence of hydrolysis degree on cellular function, nutritional content, flavor profiles, and the metabolomic implications of peptides on cellular pathways is imperative.

### 3.6. Fluorescent staining and LDH activity

The results of the combined Hoechst and actin green staining for cells in media with different concentrations of serum (0, 1, 2.5, and 10%) and also media containing 1% serum with different protein hydrolysates are presented in Fig. 5A. The results showed that serum deprivation has direct effects on the cytoskeleton, although there was no discernible effect on the nucleus other than a reduction in numbers. At 10% serum control, a high number of nuclei and abundant actin staining were observed. At 2.5% serum, there was no visible effect on the nucleus other than a reduction in cell number. However, the actin filaments densities reduced, indicating a decrease in the cell number. At 1% and serum free media conditions, the cell number reduced significantly.

Overall, the actin staining decreased along with the serum concentration, and subsequently altered the cell morphology. A similar result was reported and demonstrated that these was a decrease in actin protein staining under serum-starved circumstances (Wallenstein et al., 2010). The media with 1% serum and different protein hydrolysates, showed that the cells morphology was similar to the cell morphology in 10% serum containing media, indicating that at lower concentrations of serum, providing protein hydrolysates is critical to maintain cell performance.

The results of the LDH activity of the cells which were grown in serum-free media (0%), media containing 10% serum, 1% serum with protein hydrolysates, and 2.5% serum with protein hydrolysates are presented in Fig. 5B. Cells grown in serum-free media, showed significantly higher LDH activity compared to other conditions, indicating damage to the plasma membrane due to the starvation and lack of available serum. In contrast, cell culture at a serum concentration of 10% had the lowest LDH activity. These results suggest that the integrity of the cell membrane was compromised due to serum starvation, resulting in the release of LDH into the medium, which indicates apoptosis and necrosis. All other groups had significantly greater LDH activity compared to 10% serum control group. Cells were grown in 1% serum and protein hydrolysates had higher LDH activity compared to the 2.5% serum with protein hydrolysates. These results showed that protein hydrolysates have a critical role in maintaining the cell integrity and performance when serum is reduced or fully replaced.

### 3.7. GHG emissions

The results of the GHG emissions for different proteins, protein hydrolysates, FBS, and FBS in media at 10% and 1% concentrations, as well as 0.01 mg/mL of protein hydrolysates are presented in Table 3. Overall, the GHG emissions for algae was higher compared to other sources including FBS. During the enzymatic hydrolysis, the GHG emissions for water, electricity and enzymes were added to the protein sources. Protein hydrolysates from all the resources used in this study, had significantly higher GHG emissions, compared to FBS per FU. However, due to their low concentrations (0.01 mg/mL/10 mg/L) applications in media formulation, the GHG emission based on one liter media preparation is very small compared to serum.

**Table 3.** GHG emissions (kg CO_2_ eq.) for different biomass resources based on the literature data and calculated data in our laboratory^1^.

| Biomass | Production | Enzymatic hydrolysis process |  |  | Total (kg CO<sub>2</sub> eq./kg product) | GHG for serum in a regular media (10%) | GHG for serum (1) and protein hydrolysates (0.01 mg/mL) in reduced serum media |
| --- | --- | --- | --- | --- | --- | --- | --- |
|  |  | Enzyme | Water | Energy |  |  |  |
| Algae | 4.4 | 0.08 | 2.98 | 0.58 | 17.9 | - | $1.8 \times 10^{-4}$ |
| Pea protein | 0.94 | 0.16 | 5.96 | 1.2 | 8.26 | - | $8.26 \times 10^{-5}$ |
| Mushroom | 1.54 | 0.1 | 2.98 | 0.58 | 5.20 | - | $5.2 \times 10^{-5}$ |
| Yeast peptone | - | - | - | - | 3.2 | - | $3.2 \times 10^{-5}$ |
| FBS | 1.8 | - | - | - | 1.8 | 0.18 | 0.018 |

<sup>1</sup> Production GHG was collected from literature data: Algae (Clarens et al., 2010; Handler et al. 2012); Pea protein (Nette et al., 2016); Mushroom (American Mushroom, 2024); Yeast peptone (Carbon Cloud, 2024); FBS (Nikkhah et al., 2023). The enzymatic hydrolysis GHG was calculated based on the available literature, water, and electricity consumption during the processing. Yeast peptone was purchased directly from the provider, and no enzymatic hydrolysis was conducted in our research.

## 4. Conclusion

In this study, we assessed the impact of different concentrations of serum, both individually and in conjunction with varied concentrations of protein hydrolysates sourced from algae, mushrooms, peas, and yeast, on cell growth and viability over a short period (3 days). Reducing serum concentrations in the media, resulted in cell starvation, cell death, and significantly lower cell viability. When combined with higher concentrations of protein hydrolysates (1 and 10 mg/mL), cell apoptosis was observed in media containing 10% serum, indicating that providing higher concentrations of protein hydrolysates in conjunction with serum, will reduce the cell growth. While, lower concentrations of serum (1 and 2.5%) and protein hydrolysates (0.001, 0.01, and 0.1 mg/mL) enhanced cell growth, and cell viability, more than media containing 10% serum. Additionally, cellular fluorescence imaging analysis has shown that protein hydrolysates supplementation with reduced serum content in cell culture can improve cytoskeleton density to a similar extent as 10% serum control. GHG data showed that by applying protein hydrolysates at low concentrations, the GHG will be reduced by 90% compared to conventional media containing 10% serum.

## Data availability

Data will be made available on request.

## Ethical statement - studies in humans and animals

This work did not involve the use of human and animal subjects.

## Acknowledgment

This research was financially supported by the Agriculture and Food Research Initiative (AFRI) Sustainable Agricultural Systems program, Grant No. 2021-699012-35978 from the USDA National Institute of Food and Agriculture and Texas A&M AgriLife Research.

